# Viscoelastic Niches Shape γδ T-Cell Phenotype and Effector Function

**DOI:** 10.64898/2026.07.23.740350

**Authors:** Favour Omafuvwe Obuseh, Michelle Chang, Joshua Price, Kyle Ruark, Tania To, Leah Lourenco, Bogdan Budnik, David Mooney

## Abstract

γδ T-cells, which are predominantly enriched in epithelial tissues, have been used in cancer therapy because of their capacity for rapid cytotoxicity in an MHC-independent manner. Current paradigms are largely agnostic to the role of tissue mechanical cues in regulating γδ T-cell function. Here, we investigated the role of matrix viscoelasticity in modulating γδ T-cell migration, differentiation state, phenotype, and function. Using a tunable collagen-based gel system, we found that encapsulation in highly-elastic (slow-relaxing) matrices preserved a less differentiated phenotype, as evidenced by CD27 and CD45RA expression. Slow-relaxing matrices also increased expression of Fas and PD-1, while decreasing expression of CD11a. Despite increased PD-1 expression, these cells remained functional, as demonstrated by high levels of TNF-α and IFN-γ relative to PD-1-negative cells. Proteomic analysis revealed that γδ T-cells respond to changes in viscoelasticity through actin remodeling and shifts in metabolic machinery. Overall, when compared to non-encapsulated cells (2D culture), encapsulated γδ T-cells showed increased expression of cytotoxic programs. Functionally, cells encapsulated in slow-relaxing gels showed improved control of tumor growth in an aggressive HCT116 tumor model. Together, these findings establish matrix viscoelasticity as an important regulator of post-thymic γδ T-cell differentiation and function.

## Main Text

γδ T-cells have recently gained considerable interest in the field of cancer immunotherapy, although they have shown mixed clinical success to date. Unlike their αβ T-cell counterparts, they possess both TCR-mediated recognition and NK-like recognition mechanisms, enabling detection of metabolites such as isopentenyl pyrophosphate, aminobisphosphonates, and stress-induced ligands in a largely MHC class I–independent manner^1–4^. These interactions trigger cytotoxic responses, including TNF-α mediated apoptosis, independent of MHC presentation, providing a potential advantage in solid tumors, where MHC downregulation is common^2,4,5^.

The presence of tumor-infiltrating γδ T-cells has also been associated with improved prognosis in several solid cancers^5–9^. Together, these findings have prompted efforts to expand γδ T-cells for therapeutic application from peripheral blood, where they comprise approximately 1–5% of circulating T-cells^1,5,10^. Clinical trials evaluating these cells in hematologic malignancies have shown success, but limited efficacy has been found to date in the context of solid tumors^10–20^. While the role of cytokine combinations, including IL-15 and IL-18, in γδ T-cell cytotoxic phenotype has been studied^21,22^, little attention has been given to the physical cues that define tissue environments of these cells. A majority of clinical studies have been performed with Vδ2 T-cells, which are enriched in circulation, whereas the Vδ1 subset is more abundant in epithelial tissues^23,24^. During migration to tissues, and during residence in tissues, T-cells experience mechanical forces including shear stress, compression, and deformation to their cytoskeleton as they extravasate and navigate confined spaces^25–28^. Vδ1 T-cells are predominantly localized to tissues such as the skin, lung, and gastrointestinal tract, where they perform cytotoxic and tissue-maintaining functions^23,24,29,30^. Further, recent transcriptomic analyses have shown that γδ T-cells exhibit tissue-specific gene expression programs^22,24^, with PD-1 expression associated with tissue-resident states^12,31,32^. Despite the disparate tissue microenvironment in which γδ T-cells exist, little attention has been paid to the role of the tissue microenvironment in shaping γδ T-cell phenotype.

Here we explore the hypothesis that changes in matrix viscoelasticity shape γδ T-cell phenotype and function. Unlike stiffness, which defines a materials instantaneous response to deformation, viscoelasticity describes the time-dependent mechanical response of tissues^33,34^. In viscoelastic materials, the stress required to maintain an applied strain decreases over time, with a more rapid decline in fast-relaxing, viscous-like tissues, and a slower decline in slow-relaxing, elastic-like tissues^34,35^. Previous studies using extracellular matrix–based systems have demonstrated that viscoelasticity can regulate αβ T-cell phenotype and function, with increased elasticity promoting cytotoxic phenotypes^35–38^. Given the enrichment of γδ T-cells in epithelial tissues, which exhibit diverse viscoelastic properties^39–45^, we generated model tissue matrices using a type-1 collagen system with varying levels of viscoelasticity and utilized for γδ T-cell culture. These studies revealed that matrix viscoelasticity regulates γδ T-cell migration, phenotype, and cytotoxic function.

## Results

### Single cell RNA-sequencing reveals tissue-dependent transcriptional states in primary human γδ T-cells

To verify and extend reports that healthy human γδ T-cells in diverse tissues exhibit distinct gene expression profiles, published single-cell RNA sequencing (scRNA-seq) data were first analyzed to compare γδ T-cells found in lung, jejunum and spleen from healthy individuals. Cells originally sorted as TCRγδ^+^ by FACS were labeled as Vδ1 or Vδ2 according to *TRDV1* and *TRDV2* transcript expression, with cells exhibiting zero or equal *TRDV1* and *TRDV2* transcript expression labeled “other”. UMAP analysis revealed that the γδ T-cell populations clustered mainly by tissue source rather than Vδ1 / Vδ2 classification (Fig. 1a). Principal component analysis was then performed (Fig. 1b), revealing that γδ T-cells from the different tissues exhibited varied transcriptional profiles. The gene loadings for principal component (PC) 1, which primarily distinguishes between tissue source of γδ T-cells, exhibited enrichment of genes associated with cytoskeletal organization (KRT8, HSPB1), extracellular matrix interaction (LAIR2), and mechanosensitive signaling (F2R, TGFBR3) (Fig. 1c). PC2, which distinguished both γδ T-cell tissue of origin and Vδ1 / Vδ2 classification, reflected an orthogonal axis of proliferative and migratory state, highlighted by genes such as TK1, CKAP2L, CCR7, and GPR18, with additional contribution from immune activation-associated genes including SERPINB9 and GBP2 (Fig. 1d). To allow for comparison with circulating γδ T-cells, an additional published scRNA-seq dataset (Fig. S1a) was compared to the tissue-resident γδ T-cells dataset, using a within-dataset normalization strategy. Circulating γδ T-cells also showed significant differences in transcription factor activity from tissue-resident cells (Fig. S1b,c).

**Figure 1.**
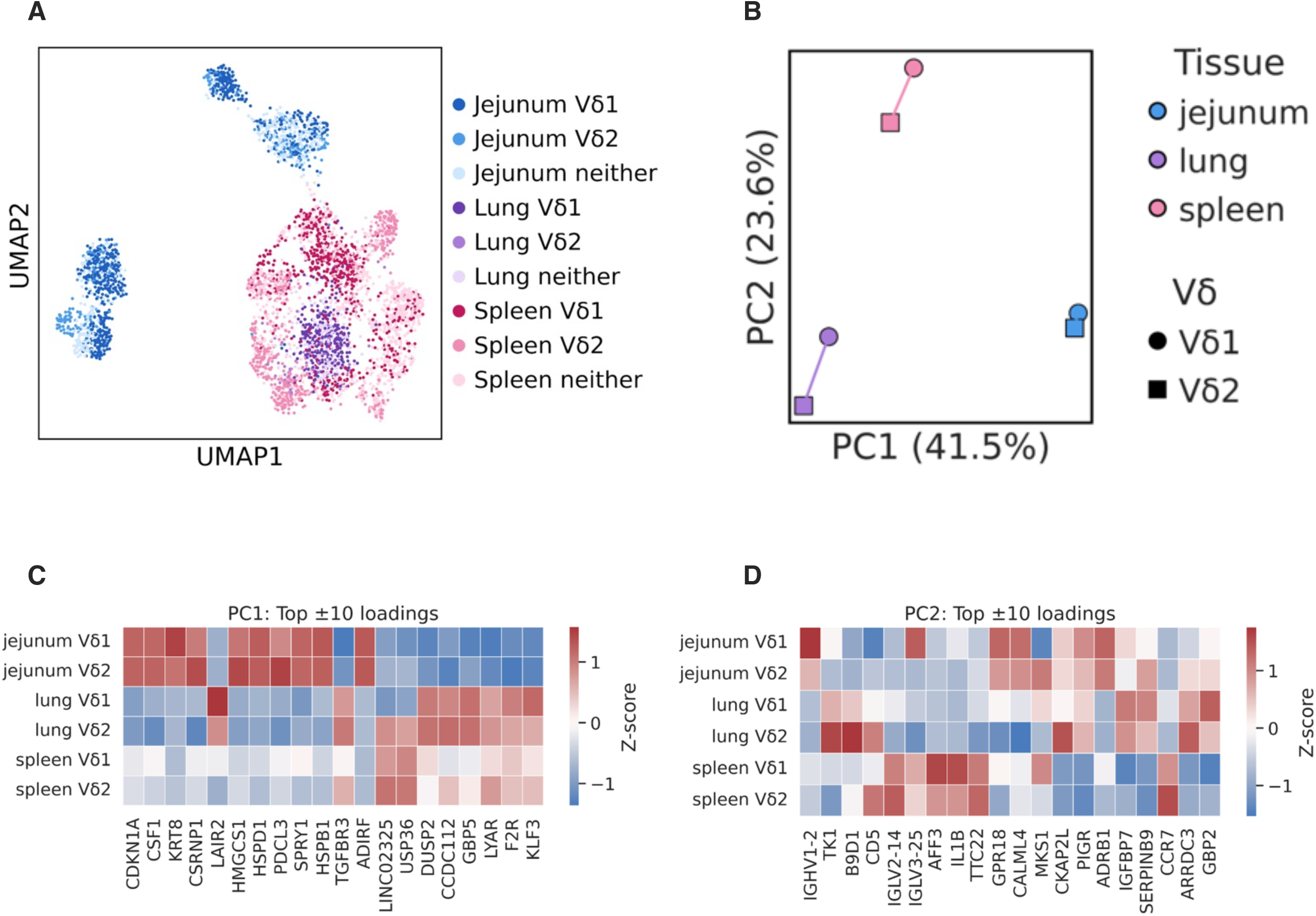
Single cell RNA-sequencing reveals tissue-dependent transcriptional states in primary human γδ T-cells. (A) UMAP of γδ T-cells from human jejunum, lung, and spleen, indicating Vδ1 and Vδ2 identities (sequencing data from^24^). (B) PCA plot of Vδ1 and Vδ2 cells from jejunum, lung, and spleen. The ten most positive and ten most negative principal component loadings for PC1 (C) and PC2 (D) from the analysis in (B), z-scored by gene. Plots represent 4 jejunum, 2 lung, and 6 spleen donors.

### Engineered collagen hydrogels enable independent tuning of viscoelasticity to model tissue mechanics

Since the initial characterization of γδ T-cells from different tissues revealed diverse profiles separated spatially by expression of genes such as KRT8, and LAIR2 which are relevant to cytoskeletal organization and extracellular matrix interaction, we hypothesized that the varying mechanical properties of different tissues contribute to shaping the cell genotype. In particular, we investigated the impact of viscoelasticity. Viscoelasticity, defined as the time-dependent response to applied stress, has been shown to significantly impact dendritic cells and conventional T-cells. T-cells actively generate cytoskeletal forces and migrate through tissue spaces, deforming their surrounding microenviroenment and experiencing time-dependent mechanical resistance. Considering that γδ T-cells are predominantly resident in epithelial tissues with distinct viscoelastic properties, we focused on this mechanical feature of the tissue microenvironment. To test our hypothesis, we utilized a collagen type I–based hydrogel system that enables independent tuning of matrix stiffness and viscoelasticity for T-cell culture. Type I collagen was modified with norbornene (Nb), which undergoes a highly selective bioorthogonal click reaction with tetrazine (Tz) moieties via an inverse electron-demand Diels–Alder reaction^35,46^. The addition of local covalent crosslinking to the Nb-modified collagen after gelation, using low-molecular-weight methyltetrazine crosslinkers, regulated matrix viscoelastic properties without altering stiffness (Fig. 2b). Characterization revealed distinct stress-relaxation profiles for the crosslinked (slow-relaxing) and non-crosslinked (fast-relaxing) conditions (Fig. 2c,d). Slow-relaxing gels corresponded to highly elastic matrices, like those observed in tissues such as the colon^42–45,47^, whereas fast-relaxing gels corresponded to highly viscoelastic matrices, as seen in lung tissue^39–41^. The overall experimental workflow for subsequent studies, following peripheral blood isolation, is shown in (Fig. 2e.)

**Figure 2.**
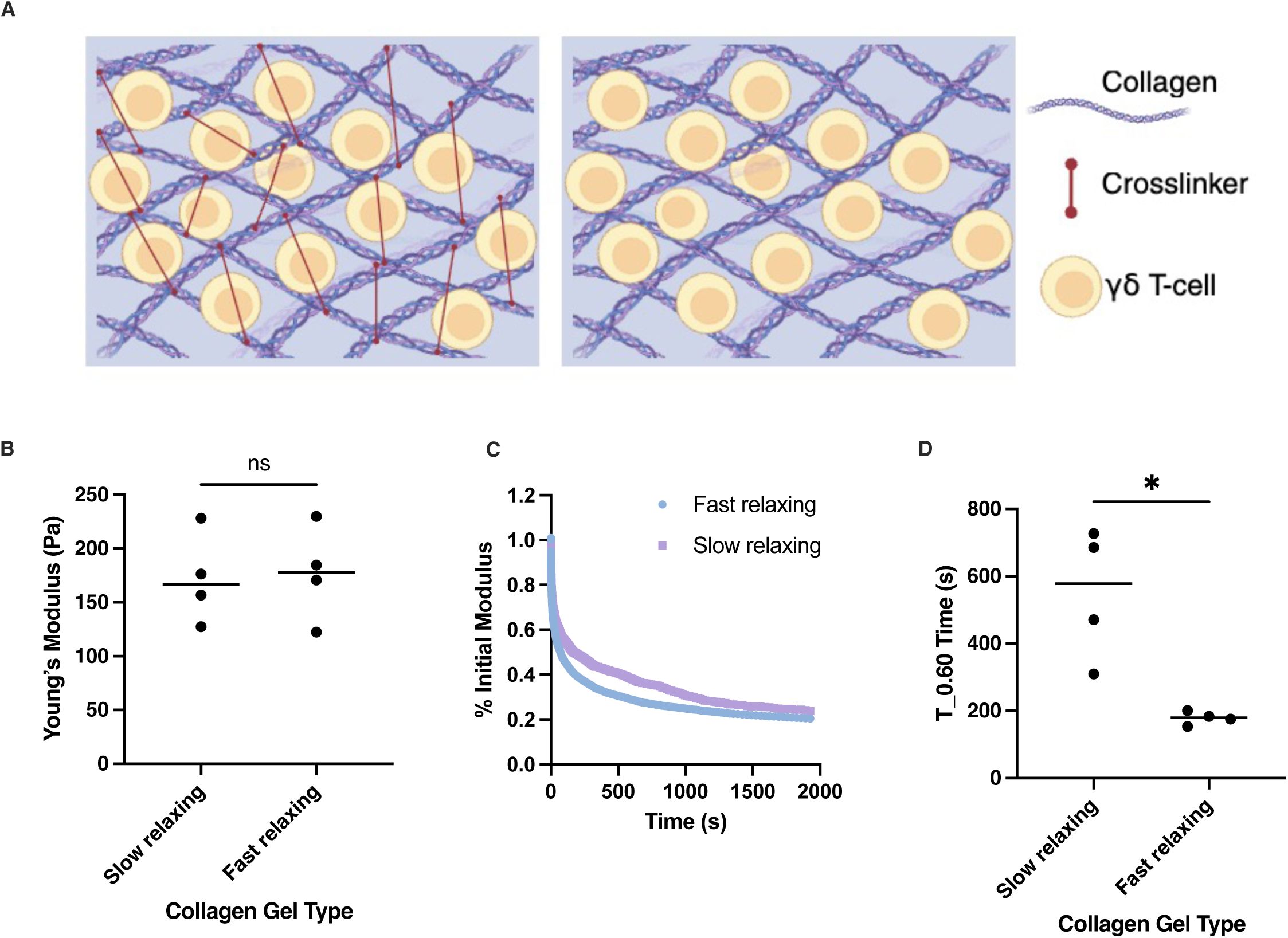
Engineered collagen hydrogels enable independent tuning of viscoelasticity to model tissue mechanics. (A) Schematic of modified collagen gels crosslinkable via norbornene–tetrazine click chemistry to allow control over viscoelasticity (B) Characterization of the shear Young’s modulus (C) Representative stress relaxation plot for slow versus fast relaxing gels. (D) Time to 60% relaxation of initial stress under constant deformation. For panel B, n = 4 technical replicates. C, n = 1, D, n= 4 technical replicates. Bar graphs show mean ± s.d. Statistical significance was determined using Mann-Whitney test. *P < 0.05, **P < 0.01, ***P < 0.001, ****P < 0.0001; ns, not significant.

### Matrix viscoelasticity modulates γδ T-cell migration

The effect of varying matrix viscoelasticity on γδ T-cell migration behavior was first studied, as migration is a fundamental aspect of T-cell function. Human-derived γδ T-cells, pre-activated for 2 days using TransAct (a non-polymeric CD3/CD28 activator), were encapsulated in hydrogels of differing viscoelasticity. Following encapsulation, cells embedded within the gels were imaged via confocal microscopy at day 1 and after 5 days of culture (Fig. 3a). For each time point, cell migration was tracked over 6-hours, with the trajectories digitized using imageJ (Fig. 3b,c). On day 1, there was no difference in mean squared displacement (MSD), a representative of the average distance the cells migrated over time, or average speed of migration speed between γδ T-cells in slow and fast-relaxing matrices (Fig. 3d-g). After 5 days, however, both MSD and average speed were greater in the fast-relaxing condition compared to the slow-relaxing condition (Fig. 3h-k), suggesting an impact of physical encapsulation in distinct environments on propensity for migration.

**Figure 3.**
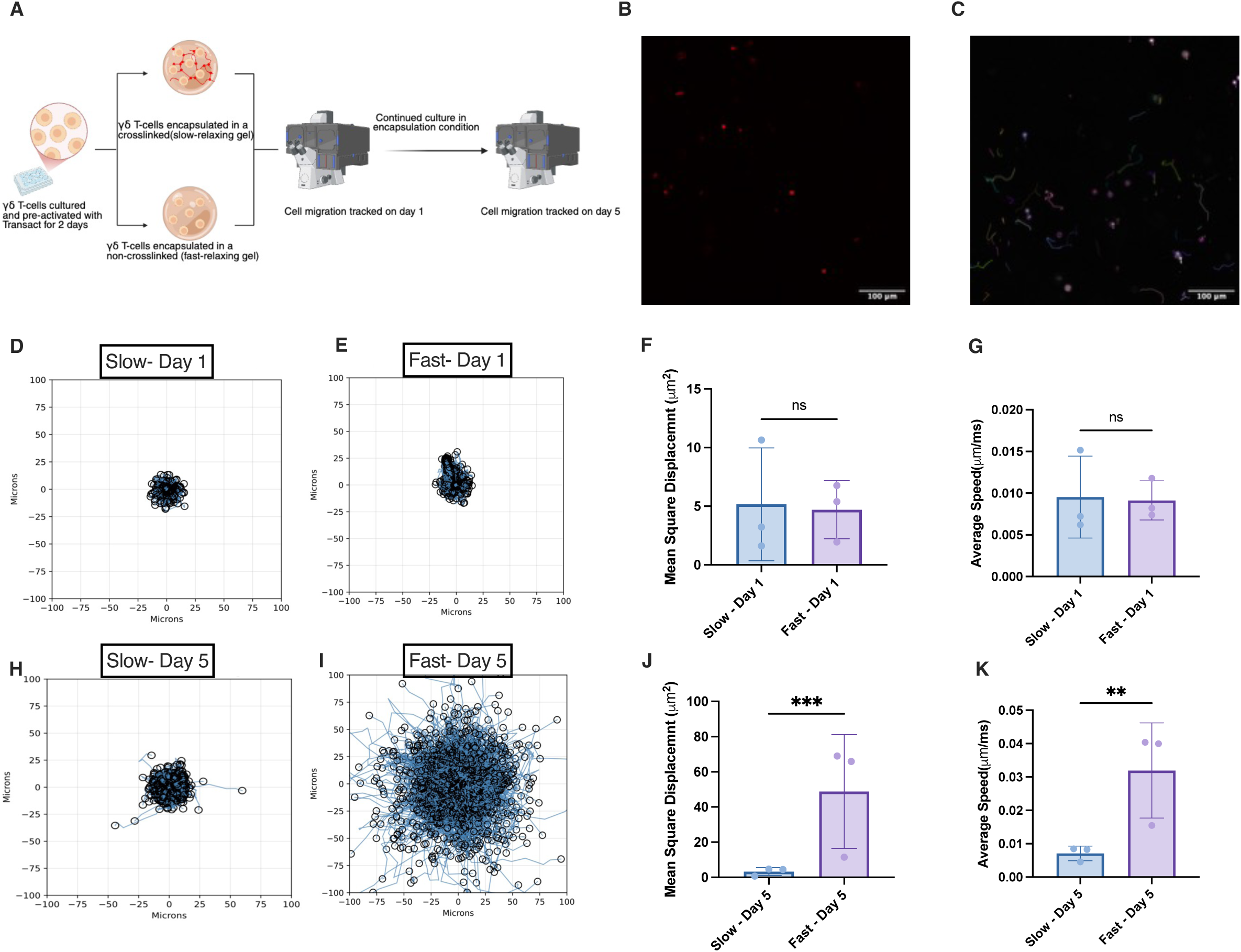
Matrix viscoelasticity modulates γδ T-cell migration. (A) Schematic of the experimental workflow for cell migration quantification. (B) Confocal images of fluorescently labeled γδ T-cells within the gel matrix. Scale bar: 100 µm. (C) Representative ImageJ tracking overlay. Migration tracks for γδ T-cells in slow-relaxing (D) and fast-relaxing (E) gels on day 1 of encapsulation.Average mean squared displacement (F) and average speed (G) on day 1 for cells encapsulated in slow versus fast-relaxing matrices. Representative migration tracks for γδ T-cells in slow-relaxing (H) and fast-relaxing (I) gels five days post-encapsulation. Average mean squared displacement (J) and average speed (K) on day 5 for cells encapsulated in slow-versus fast-relaxing matrices. For panels F, G, J, and K, n = 3 donors. For panels D, E, H, and I, n = 1 donor. Bar graphs show mean ± s.d. Statistical significance was determined using a ratio paired t-test. *P < 0.05, **P < 0.01, ***P < 0.001, ****P < 0.0001; ns, not significant.

### Matrix elasticity reprograms γδ T-cells toward a tissue-adapted cytotoxic phenotype

To assess phenotypic changes that could accompany the migration differences, cells were again encapsulated and subsequently analyzed. In these studies, cells were also maintained in standard 2D cell culture as a control.

Cells were retrieved from the different matrices on day 5 for various analysis (Fig. 4a) There were no significant differences in viability across all conditions (Fig. S2a). Cell differentiation, as based on CD27 and CD45RA classification, was decreased overall in γδ T-cells encapsulated in the more elastic (slow-relaxing) gels (Fig. 4b). When stratified into the two major subsets, a significant difference in the Vδ1/Vδ2 ratio was observed compared to non-encapsulated cells, with a higher ratio in the slow-relaxing condition (Fig. S2b).Within the Vδ1⁺ γδ T-cell population, studies with cells from multiple donors revealed the largest fraction of cells were CD27⁺CD45RA⁺, suggesting maintenance of a large pool of naïve cells. The fraction of these cells was largest in the slow relaxing gels, with a decrease in fast relaxing gels, and the lowest fraction in 2D culture. No statistically significant differences were observed in the CD27⁺CD45RA⁻, CD27⁻CD45RA⁻, or CD27⁻CD45RA⁺ populations (Fig. 4c). In the Vδ2 subset, the fraction of CD27⁺CD45RA⁺ cells was significantly diminished, but encapsulation in slow-relaxing gels best preserved this population, while also demonstrating the largest fraction of CD27⁺CD45RA⁻ cells, which is associated with central memory related differentiation. In contrast, encapsulation in more rapidly relaxing gels was associated with a greater CD27⁻CD45RA⁻ population, which is associated with effector memory populations, and a trend of increased CD27^-^CD45RA^+^ cell population (Fig. 4d).

**Figure 4.**
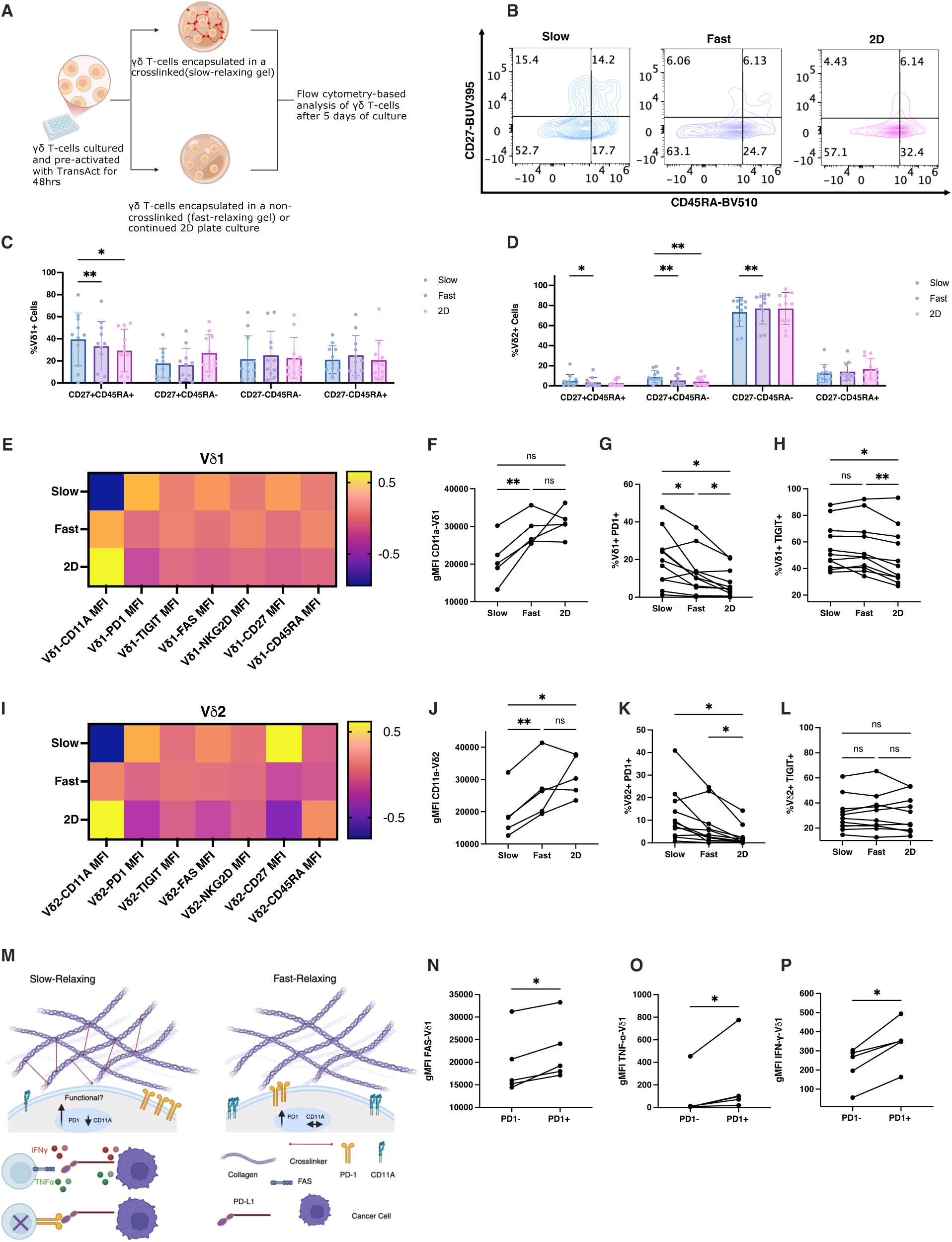
Matrix elasticity reprograms γδ T-cells toward a tissue-adapted cytotoxic phenotype. (A) Schematic of the general workflow following isolation of γδ T-cells from peripheral blood. (B) Representative flow plot of CD27 and CD45RA percentages for γδ T-cells after encapsulation in slow-relaxing, fast-relaxing, or unencapsulated (2D) conditions.(C) CD27 and CD45RA percentages for Vδ1 γδ T-cells after encapsulation in slow-relaxing, fast-relaxing, or unencapsulated (2D) conditions. (D) CD27 and CD45RA percentages for Vδ2 γδ T-cells after encapsulation in slow-relaxing, fast-relaxing, or unencapsulated (2D) conditions. (E) Heat maps with column-wise z-scored, log-transformed geometric mean fluorescence intensity (gMFI) of CD11a, PD-1, TIGIT, FAS, NKG2D, CD27, and CD45RA for Vδ1. (F) Log-transformed geometric mean fluorescence intensity (gMFI) of CD11a expression in Vδ1⁺ cells. (G) Percentage of Vδ1⁺ PD-1⁺ cells after encapsulation in slow-relaxing, fast-relaxing, or unencapsulated (2D) conditions. (H) Percentage of Vδ1⁺ TIGIT⁺ cells after encapsulation in slow-relaxing, fast-relaxing, or unencapsulated (2D) conditions. (I) Heat maps with column-wise z-scored, log-transformed geometric mean fluorescence intensity (gMFI) of CD11a, PD-1, TIGIT, FAS, NKG2D, CD27, and CD45RA for Vδ2 (J) Log-transformed geometric mean fluorescence intensity (gMFI) of CD11a expression in Vδ2⁺ cells. (K) Percentage of Vδ2⁺ PD-1⁺ cells after encapsulation in slow-relaxing, fast-relaxing, or unencapsulated (2D) conditions. (L) Percentage of Vδ2⁺ TIGIT⁺ cells after encapsulation in slow-relaxing, fast-relaxing, or unencapsulated (2D) conditions. (M) Schematic of current findings and expected outcome with increased PD-1 expression. (N–P) Expression of FAS, TNFα, and IFNγ in Vδ1⁺ PD-1⁺ versus PD-1⁻ cells, respectively. For panels C and D, n = 11 donors. For panels E and I, n = 5–11 donors. For panels F, J, and N-P, n = 5 donors. For panels G, H, K, and L, n = 11 donors. Bar graphs show mean ± s.d. Statistical significance was determined using ordinary two-way ANOVA for C-D. One-way ANOVA with Tukey’s multiple comparisons test for F-H, and J-L, Statistical significance for N-P was determined using a ratio paired t-test. *P < 0.05, **P < 0.01, ***P < 0.001, ****P < 0.0001; ns, not significant.

Further phenotypic analysis was next performed, and geometric mean fluorescence intensity (gMFI) analysis showed increased expression of CD11a, PD-1, Fas, and TIGIT in both Vδ1 and Vδ2 cells encapsulated in slow relaxing gels (Fig. 4e,f). When examining data in a donor specific manner, encapsulation of Vδ1 cells in slow-relaxing gels resulted in a significant decrease in CD11a expression compared to fast-relaxing and non-encapsulated (2D) conditions (Fig. 4g). However, there was a greater percentage of PD-1⁺ cells in slow relaxing gels, with a further decrease when comparing fast-relaxing and 2D conditions (Fig. 4h). No difference in TIGIT⁺ cell frequency was observed as a function of gel viscoelasticity; however, both slow and fast-relaxing conditions showed increased TIGIT⁺ frequency relative to the 2D condition (Fig. 4i). For Vδ2 cells, encapsulation in slow-relaxing gels resulted in a significant decrease in CD11a expression relative to fast-relaxing and 2D conditions (Fig. 4j). A statistically significant increase in PD-1⁺ cells was observed with encapsulation, independent of viscoelasticity (Fig. 4k). No differences in TIGIT⁺ cell frequency were observed across conditions (Fig. 4l). Given the observed increase in PD-1⁺ cells in encapsulated conditions, we examined whether PD-1 expression was associated with functional changes (Fig. 4m). Extracellular and intracellular staining after stimulation with cancer cells revealed that PD-1⁺ cells exhibited higher expression of Fas, TNFα, and IFNγ compared to PD-1⁻ cells (Fig. 4n-p).

### Differential matrix viscoelasticity drives actin–associated proteomic remodeling in γδ T-cells

To provide a broad assessment of the impact of matrix viscoelasticity and physical encapsulation on γδ T-cells, Vδ1 and Vδ2 T-cells derived from slow-relaxing, fast-relaxing, and 2D conditions were FACS sorted and analyzed by LC–MS.

Encapsulation of Vδ1 T-cells in slow-relaxing gels led to increased abundance of proteins associated with metabolic processes and mitochondrial function, including SFXN1, MECR, GLS, GCDH, GYS1, PGM1. (Fig. 5a,b). Proteins related to protein synthesis and processing and transcriptional and RNA regulatory pathways, cellular organization, and homeostasis, were also enriched in the cells from the slow-relaxing condition, (Dataset 2). Compared to slow-relaxing matrices, the Vδ1 T-cells from the fast-relaxing matrices were enriched for proteins related to cytoskeletal reorganization and actin dynamics, including ARPC2, RAC1, BRK1, RDX, and G3BP2. Immune-associated proteins (GZMA, CCL5, PSMB10)(Fig. 5a,b), proteins involved in vesicular trafficking and membrane organization, transcriptional and RNA processing pathways, cell cycle regulation and DNA replication/repair were also increased in cells from fast-relaxing gels (Dataset 2). Gene ontology (GO) pathway analysis of differentially expressed proteins revealed enrichment of pathways associated with actin filament organization in Vδ1 T-cells (Fig. 5e).

**Figure 5.**
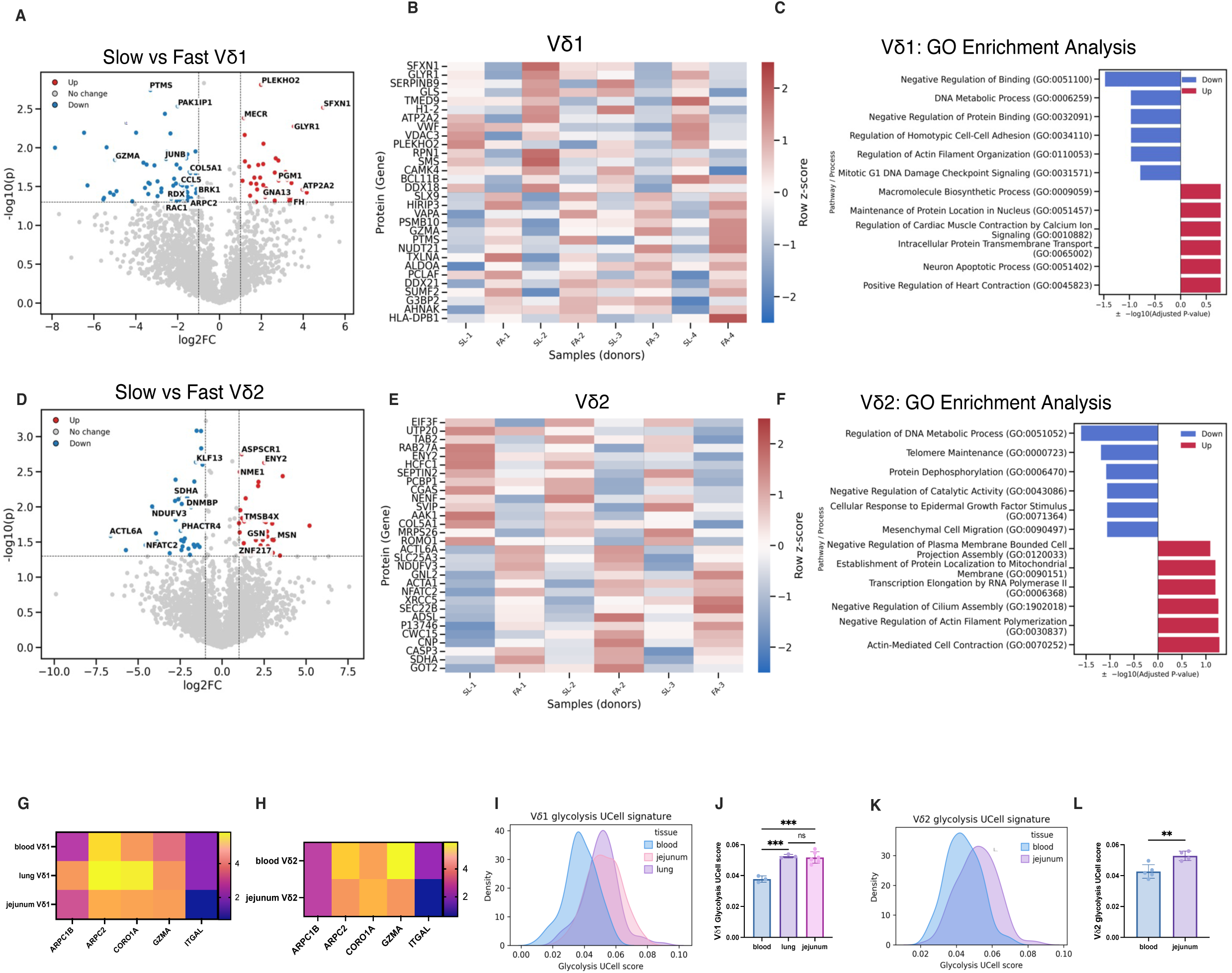
DiKerential matrix viscoelasticity drives actin–associated proteomic remodeling in γδ T-cells. (A) Volcano plot comparing FACS-sorted Vδ1 cells cultured in slow versus fast-relaxing matrix conditions. (B) Top 30 proteins ranked by composite score (log₂ fold change × –log₁₀ p-value) for Vδ1 cells cultured in slow versus fast-relaxing matrix conditions. C) Enrichr over-representation analysis (ORA) of Gene Ontology (GO) biological processes for Vδ1 cells cultured in slow versus fast-relaxing matrix conditions. (D) Volcano plot comparing FACS-sorted Vδ2 cells cultured in slow versus fast-relaxing matrix conditions. (E) Top 30 proteins ranked by composite score (log₂ fold change × –log₁₀ p-value) for Vδ2 cells cultured in slow versus fast-relaxing matrix conditions.(F) Enrichr ORA of GO biological processes for Vδ2 cells cultured in slow versus fast-relaxing matrix conditions. (G) Relative expression of genes associated with γδ T-cell function in blood, jejunum, and lung-Vδ1 cells (z-scored across all genes for each donor.(H) Relative expression of genes associated with γδ T-cell function in blood and jejunum-Vδ2 cells (z-scored across all genes for each donor). Kernel density estimate plots (I) and donor-bulked averages (J) of glycolysis UCell signature scores in Vδ1 γδ T-cells from blood, jejunum, and lung. Kernel density estimate plots (K) and donor-bulked averages (L) of glycolysis UCell signature scores in Vδ2 γδ T-cells from blood and jejunum For panels A-C, n = 4 donors; for panels D-F, n = 3 donors. (G,I, J), n= 3 blood, 3 lung, and 6 jejunum donors. (H,K,L) n= 5 blood and 4 jejunum donors. Bar graphs show mean ± s.d. Statistical significance in A,D, was determined using a paired t-test. Statistical comparisons between tissues in (J) was performed using Prism’s ordinary one-way ANOVA with Tukey’s post hoc multiple comparisons test. An unpaired t-test was carried out for (L). *P < 0.05, **P < 0.01, ***P < 0.001; ns, not significant.

Encapsulation of Vδ2 T-cells in slow-relaxing gels increased the abundance of proteins associated with cytoskeletal organization and actin dynamics, including MSN, GSN, TMSB4X, and LUZP1 (Fig. 5d,e). Proteins involved in metabolic and mitochondrial function (MRPS26, ECI1, ROMO1, TOMM5), transcriptional regulation and RNA processing, protein synthesis, vesicular trafficking, and cellular signaling pathways were also increased (Dataset 2). Compared to slow-relaxing matrices, the Vδ2 T-cells from the fast-relaxing gels were enriched for proteins associated with metabolic and mitochondrial function (SDHA, NDUFV3, SLC25A3), cytoskeletal organization and membrane dynamics (IQGAP1, DNMBP, PHACTR4) (Fig. 5d, e). Proteins related to cell cycle and DNA repair, RNA processing, vesicular trafficking were also increased in these cells. Additional increases were observed in proteins associated with cellular signaling, (Dataset 2). Gene ontology (GO) pathway analysis of differentially expressed proteins also revealed enrichment of pathways associated with actin filament organization in Vδ2 T-cells (Fig. 5f).

To assess the impact of 3D encapsulation, cells from fast-relaxing and 2D conditions were also compared, revealing enrichment in Vδ1 T-cells, proteins associated with immune activation and cytotoxic function (GZMA, ZAP70, IKBKG, EOMES), actin remodeling and cytoskeletal reorganization (ARPC2, AHNAK, ANXA2, HCLS1, BIN1, S100A4) (Fig. S3a). Proteins associated with metabolic pathways, vesicular trafficking, transcriptional and RNA processing pathways, and cell cycle regulation were also increased. Conversely, proteins decreased in fast-relaxing gels compared to 2D culture included those associated with metabolic function (UQCRC1, LDHB, NDUFA6), cytoskeletal organization and membrane dynamics (TPM1, STOM, AKAP13, MICALL1), (Fig.S3a, Dataset 2). Encapsulated Vδ2 T-cells similarly increased proteins associated with actin architecture organization (FSCN1, MYO1G, FYB1, ESYT1), and metabolic and mitochondrial function (COQ7, CHCHD2, MRPS23, MRPS6, PNPT1) (Fig. S3b). Proteins involved in RNA processing and translation, vesicular trafficking, and cellular signaling pathways were also enriched. Proteins associated with actin filament stabilization and vesicular trafficking (TPM3 and BLOC1) were reduced in encapsulated cells, as well as metabolic regulation relevant proteins (SLC7A5, ENTPD1) (Fig.S3b, Dataset 2).

Motivated by the impact of viscoelasticity on actin and metabolism associated pathways, we next examined published scRNA-seq datasets to determine whether actin-related genes varied in cells across tissues. Within-donor Z-scored expression values of individual gene expression in blood, jejunum, and lung γδ T-cells were compared. In Vδ1 T-cells, the actin-associated genes ARPC1B, ARPC2, CORO1A, and ITGAL were differentially expressed across tissues, with increased expression observed in lung compared to other tissues. GZMA expression was also higher in both lung and jejunum relative to blood-Vδ1 T-cells (Fig. 5g). For Vδ2 T-cells, due to limited cell numbers from lung, comparisons were restricted to blood and jejunum. Relative ARPC2, GZMA, and ITGAL, expression were higher in blood compared to jejunum Vδ2 T-cells, whereas CORO1A was increased in jejunum Vδ2 T-cells (Fig. 5h). Circulating γδ T-cells also showed differences in energy expenditure signatures relative to tissue-resident γδ T-cells, with both Vδ1 and Vδ2 subsets showing higher expression of glycolysis-related genes in both lung and jejunum tissues compared to blood (Fig. 5i-l).

### Matrix viscoelasticity and elasticity differentially polarize γδ T-cell effector function

To evaluate the cytotoxic potential of cells derived from different matrices, intracellular cytokine staining was performed. In the Vδ1 T-cell population, cells derived from slow-relaxing matrices showed a significantly increased percentage of IFNγ⁺ and TNFα⁺ cells, whereas granzyme B expression appeared increased in the fast-relaxing condition (Fig. 6a–c). In the Vδ2 T-cell population, no significant difference in IFNγ⁺ cells were observed; however, TNFα⁺ cells were significantly increased in the slow-relaxing condition, while granzyme B⁺ cells were significantly increased in the fast-relaxing condition (Fig. 6d–f). An adoptive cell transfer model using a HCT116 human colorectal xenograft tumor, (Fig 6g), showed that cells derived from slow-relaxing matrices provided more control over tumor burden when compared to cells derived from fast-relaxing and unencapsulated 2D controls. (Fig. 6h).

**Figure 6.**
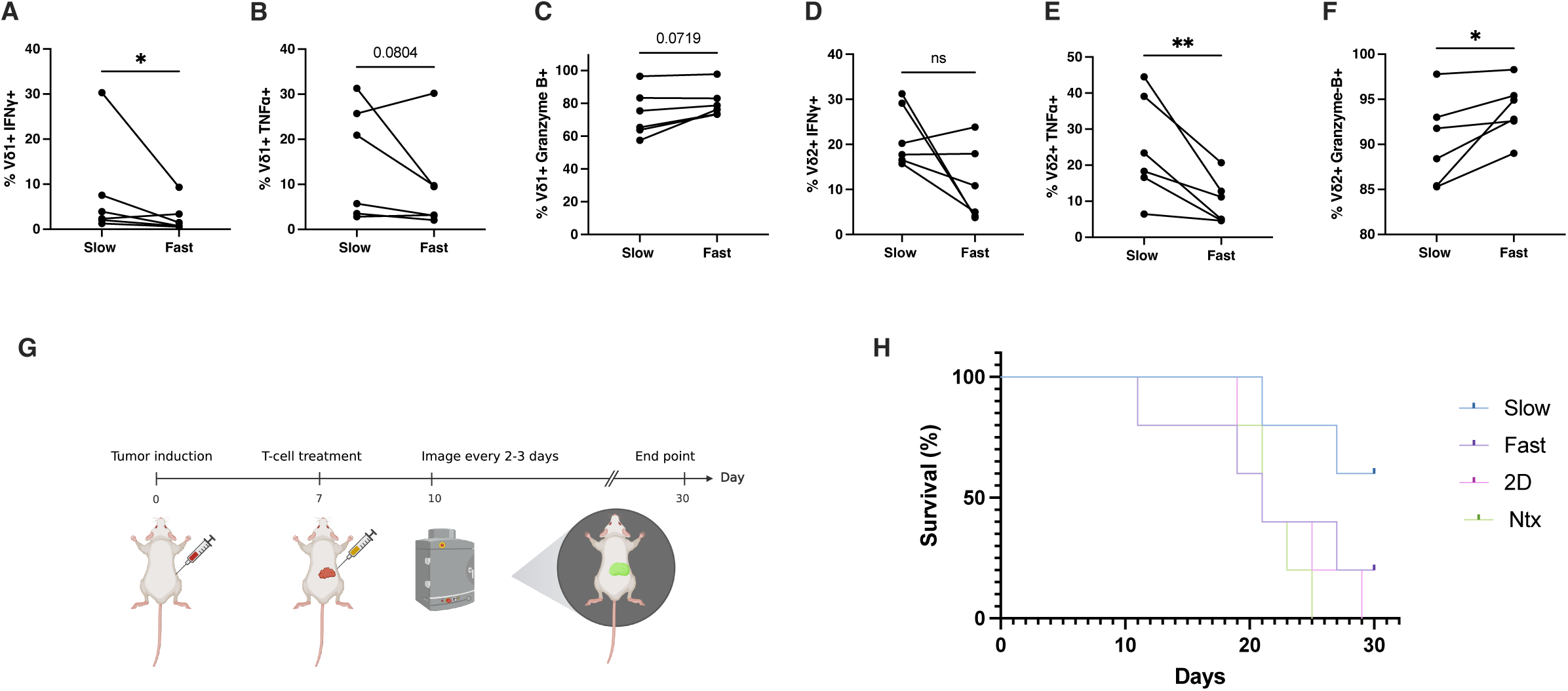
Matrix viscoelasticity and elasticity diKerentially polarize γδ T-cell eKector function. (A) Percentage of IFNγ⁺ Vδ1 T-cells after encapsulation in slow-relaxing and fast-relaxing gels. (B) Percentage of TNFα⁺ Vδ1 T-cells after encapsulation in slow-relaxing and fast-relaxing gels. (C) Percentage of granzyme B⁺ Vδ1 T-cells after encapsulation in slow-relaxing and fast-relaxing gels.(D) Percentage of IFNγ⁺ Vδ2⁺ T-cells after encapsulation in slow-relaxing and fast-relaxing gels. (E) Percentage of TNFα⁺ Vδ2 T-cells after encapsulation in slow-relaxing and fast-relaxing gels. (F) Percentage of granzyme B⁺ Vδ2 T-cells after encapsulation in slow-relaxing and fast-relaxing gels.(G) Schematic of in vivo tumor control study.(H) Kaplan Meier Survival Curve for slow-relaxing, fast-relaxing, 2D, and non-treated (NTx) conditions. For panels A–F, n = 5 donors; for panel H, n= 4–5 donors; Statistical significance for A-F was determined using a ratio paired t-test. *P < 0.05, **P < 0.01, ***P < 0.001, ****P < 0.0001; ns, not significant.

## Discussion

A biomaterial system with tunable viscoelasticity was utilized to study how changes in matrix viscoelasticity influenced migration, differentiation state, and phenotype of γδ T-cells. Widespread proteomic remodeling was observed in response to changes in viscoelasticity, with enrichment of pathways related to actin organization. In addition, the viscoelasticity of the microenvironment modulated the cytotoxic function of γδ T-cells, establishing matrix viscoelasticity as a key regulator of γδ T-cell state and function.

A collagen-based biomaterial system demonstrated an impact of changing viscoelasticity on γδ T-cell migration, differentiation state and phenotype. Covalently crosslinking Nb-modified collagen using methyltetrazine crosslinkers allowed for control over stress relaxation without changing stiffness. Encapsulation in slow-relaxing matrices, which model highly elastic tissues such as the colon, resulted in reduced migration compared to fast-relaxing matrices. Phenotypic analysis based on CD27 and CD45RA expression^23,48^, revealed reduced differentiation from the CD27⁺CD45RA⁺ state in both Vδ1 and Vδ2 T-cells under slow-relaxing conditions. In contrast to αβ T-cells, where elastic matrices have been reported to promote differentiation toward more experienced phenotypes^35^, γδ T-cells exhibited an opposing response, suggesting fundamental differences in how these subsets interpret mechanical cues. Given that γδ T-cells are predominantly resident in epithelial tissues^1,2,9,23^, this divergence may reflect adaptation to tissue-specific environments. Consistent with this, increased Fas expression, PD-1 expression and decreased CD11a expression, in the absence of chronic activation, in slow-relaxing conditions indicate a shift toward a tissue-associated phenotype characterized by reduced migratory capacity and increased cytotoxic potential^49–51^. For αβ T-cells, reduced expression of molecules involved in tissue egress is associated with tissue resident biology^49,50,52^. For example, CD8+ tissue resident memory cells of the lung were shown to initially express high levels of CD11a, but rapidly lose expression of this integrin with time spent in the lung tissue^51^. While PD-1 is typically associated with exhaustion in conventional αβ T cells, it was observed that PD-1⁺ γδ T-cells showed increased expression of Fas, IFNγ, and TNFα compared to PD-1⁻ cells, consistent with recent findings that PD-1 expression can coincide with a functional, tissue-adapted state^31^. Previous work has shown that PD-1 can exert inhibitory effects independently of its canonical ITIM and ITSM motifs, which mediate SHP2 recruitment, suggesting a potential mechanosensitive role^53^. In that context, PD-1 engagement was shown to inhibit actin remodeling at the immunological synapse of αβ T cells through a signaling-independent mechanism. In contrast, our data indicate that in γδ T cells, PD-1 expression is associated with functional competence and tissue adaptation. These findings suggest that, for both γδ T-cell subsets, a slow-relaxing microenvironment induces a tissue-adapted phenotypic state characterized by reduced differentiation from naïve-like populations, decreased expression of migration-associated markers, and increased expression of cytotoxicity-associated modules which may be relevant to maintain tissue homeostasis.

Widespread proteomic remodeling was found in response to changes in viscoelasticity, including alteration of pathways related to actin organization. Vδ1 T-cells encapsulated in slow-relaxing gels, as compared to fast-relaxing gels, exhibited decreased expression of proteins associated with lamellipodia formation and actin organization, including ARPC2, RAC1, BRK1, and RDX^28,54–56^. In contrast, proteins associated with metabolic processes and mitochondrial function, including GLS, PGM1, and MECR, were increased in slow-relaxing conditions, consistent with enrichment of metabolic pathways identified by Gene Ontology analysis. These findings suggest a shift toward a metabolically active state^57^, while cells in fast-relaxing matrices retained features associated with migration and cytotoxic function, including increased expression of GZMA, CCL5, and PSMB10^22,24^. In contrast, Vδ2 T-cells exhibited a different response to viscoelastic cues. Cells from slow-relaxing matrices showed increased abundance of proteins associated with actin remodeling, including MSN, GSN, and TMSB4X, whereas cells from fast-relaxing matrices were enriched for proteins associated with mitochondrial metabolism and cytoskeletal stability. These differences in response between Vδ1 and Vδ2 T-cells may reflect intrinsic differences in baseline tissue distribution, as Vδ1 T-cells are predominantly tissue-resident, whereas Vδ2 T-cells are more commonly found in circulation^23,24,58,59^.

Comparison to unencapsulated cells further highlighted the effects of 3D matrix engagement. Vδ1 T-cells cultured in fast-relaxing matrices exhibited increased expression of proteins associated with cytotoxic function and signaling, including GZMA, ZAP70, IKBKG, and EOMES, whereas 2D-cultured cells displayed a comparatively limited cytotoxic profile. In Vδ2 T-cells, encapsulation was associated with coordinated changes in cytoskeletal and metabolic programs, with enrichment of proteins linked to actin remodeling and mitochondrial function relative to 2D conditions. These findings were supported by the RNA seq analysis of Vδ1 and Vδ2 T-cells from lung, jejunum and blood. Cells from lung, which can be characterized as a fast-relaxing tissue^39–41^, showed increased expression of ARPC2 and CORO1A, relative to cells from jejunum, a slow-relaxing tissue^42–45,47^, and blood a viscous non-Newtonian tissue^60,61^. ITGAL which is the gene responsible for CD11a expression was also reduced in jejunum as compared to the other tissues, consistent with the cell culture data generated here. These findings are also consistent with the known roles of the cytoskeleton in controlling how cells respond to their environment, mediating signal transduction through regulators such as the Arp2/3 complex (including ARPC2), RAC1, and BRK1, as well as actin modulators including GSN and MSN, which control actin nucleation in the context of migration and synapse formation^26,28,54–56,62,63^. In addition, cytoskeletal organization is linked to mitochondrial function, influencing cellular metabolism, activation state, and cytotoxic function^25,64–67^. Consistent with this framework, our data demonstrate that modulation of viscoelasticity is accompanied by coordinated changes in actin-associated and metabolic protein networks.

Slow-relaxing hydrogels promoted PD-1 and Fas expression, together with upregulation of activation and cytotoxicity-associated markers such as TNFα and IFNγ, while downregulating CD11a, consistent with a shift toward a tissue-retentive phenotype. Functionally, γδ T-cells from slow-relaxing matrices better controlled the growth of HCT116 tumor cells than cells cultured in fast-relaxing or 2D conditions, underscoring the ability of mechanical conditioning to enhance cytotoxic capacity. Although fast-relaxing conditions showed a higher proportion of granzyme B⁺ cells, the in vivo results suggest that additional factors contribute to tumor control. One potential explanation is that cells from slow-relaxing matrices are conditioned to function in low-adhesion, high-resistance environments, favoring mechanisms such as death receptor signaling through Fas–FasL interactions or secretion of cytotoxic factors, rather than direct immune synapse–mediated killing^35,68^. Tissue-resident memory T-cells in epithelial tissues are also known to exhibit a highly cytotoxic state and produce elevated levels of inflammatory cytokines, including IFNγ and TNFα ^49^. Together, these findings indicate that culture in highly elastic matrices induces adaptations in γδ T-cells that recapitulate features of tissue-associated states while enhancing tumor-directed cytotoxic function.

The role of matrix viscoelasticity in shaping γδ T-cell phenotype and function has in the past been largely unexplored. Here, we show that viscoelasticity, a core mechanical feature distinguishing tissues, regulates γδ T-cell migration, phenotype, cytotoxic potential, and expression of markers associated with tissue adaptation. These findings support a shift away from conventional 2D expansion systems toward mechanically tunable 3D environments for γδ T-cell culture. More broadly, our results highlight the potential importance of mechanical cues in T-cell activation, whereby physical cues from the microenvironment contribute to functional programming. Aligning engineered physical cues with known target tissue-associated transcriptional programs could inform manufacturing strategies aimed at improving the functional potency of γδ T-cells for adoptive therapies.

## Materials and Methods

### Single-cell RNA sequencing data analysis

Processed single-cell RNA sequencing (scRNA-seq) data of human γδ T-cells from primary spleen, jejunal epithelium, and lung were downloaded from https://cellxgene.cziscience.com/collections/ec691f5f-0aac-433c-8f78-e7f4b85a05e0 and are available at NCBI GEO with accession number GSE263248. Processed scRNA-seq data of human γδ T-cells from primary peripheral blood mononuclear cells were downloaded from https://cellxgene.cziscience.com/collections/b0cf0afa-ec40-4d65-b570-ed4ceacc6813 and are available at NCBI GEO with accession number GSE164378. γδ T-cells were labeled by surface protein expression of γδ T-cells receptor according to flow cytometry measurements in both studies.

Gene expression analysis was performed in Python (version 3.12.13) with the scanpy package (version 1.12.1). γδ T-cells with greater *TRDV1* transcript expression as compared to *TRDV2* were computationally labeled Vδ1, those with greater TRDV2 transcript expression were labeled Vδ2, and those with equal or no *TRDV1* and *TRDV2* expression were labeled “neither” and excluded from downstream analysis. Expression values were normalized and log1p transformed with the natural log. All analyses were performed with a minimum single donor cell count of 25 cells for each cell type included in the analysis. Neighborhood graphs and Uniform Manifold Approximation and Projection (UMAP) visualizations were generated with scanpy. Principal component analysis (PCA) was computed in Python’s scikit-learn package (version 1.6.1).

Transcription factor activity analysis was performed using Python’s decoupler package (version 2.1.6) with CollecTRI-derived regulons, enabling a gene set based measure that is less sensitive to cross-dataset technical variation than individual gene expression. Glycolysis gene set activity was quantified with the UCell package in R (version 2.14.0) by computing UCell scores on individual cells with the HALLMARK_GLYCOLYSIS gene set obtained from MSigDB (systematic name M5937), likewise enabling cross-dataset comparison that is robust to differences in sequencing depth and technical variation. Cross-tissue glycolysis UCell score statistical comparison was conducted in GraphPad Prism 10 using one-way ANOVA with Tukey’s post-hoc multiple comparisons tests on donor glycolysis UCell score means.

### Material synthesis

Rat Tail Collagen Type I (Corning, 354236) was functionalized with 5-norbornene-2-acetic acid succinimidyl ester (Nb-NHS) (Sigma Aldrich, 776173) at a ratio of 0.1 g Nb-NHS:1 g collagen. Rat tail collagen was first neutralized with NaOH and buffered with 10x Dulbecco’s phosphate-buffered saline to a concentration of 2 mg/ml. Next, Nb-NHS was dissolved in dimethylsulfoxide (DMSO) to an initial concentration of 2 mg/ml and diluted 10-fold in 1x PBS, after which the Nb-NHS solution was added to the neutralized collagen, resulting in 1 mg ml−1 final collagen concentration. The reaction proceeded for 4 hrs at 4°C under rigorous stirring to delay collagen gelation, after which 0.1 N acetic acid was added to quench the reaction and re-acidify the collagen solution. The product, Norbornene-modified collagen (Col-Nb) was dialysed in 0.025 N acetic acid for 3 d, filtered with a 0.45 μm filter and then lyophilized for future use.

### Human γδ T-cell isolation, activation, and culture

#### Human γδ T-cell isolation

Human Peripheral blood mononuclear cells (PBMCs) were obtained from Brigham and Women’s Hospital as-de-identified apheresis collars which were processed by density-gradient centrifugation using Ficoll-Paque PLUS. A fraction of fresh PBMCs was characterized by flow cytometry (see Flow Cytometry) and the remainder cryopreserved in 90% FBS / 10% DMSO. PBMCs were enriched for γδ T cells by negative selection using the Human γδ T Cell Isolation Kit (Miltenyi Biotec, 130-092-892) following the manufacturer’s instructions.

#### Human γδ T-cell culture and activation

Enriched γδ T-cells were resuspended in RPMI-1640 (Lonza, BE12-702F), supplemented with 10% heat-inactivated fetal bovine serum (Gibco,10-082-147), 1% penicillin–streptomycin, 20 mM HEPES (Gibco, 15630080), and 1x non-essential amino acids (Fisher, 11140-050), 1x sodium pyruvate (Gibco, 11360070), 55µM beta-mercaptoethanol. Cytokines were added at the following final concentrations unless otherwise indicated: 50 ng mL-recombinant human IL-18 (R&D systems, 9124-IL-050); 70 ng mL-recombinant human IL-15 (Fisher Scientific, 10773-078). For initial T-cell activation, TransAct (Miltenyi Biotec, 130-111-160) was added at a 1:100 (v/v) ratio as per manufacture’s instruction. Culture was maintained at 37°C, 5% CO₂.

#### Cell encapsulation and crosslinking

After 2 days of activation, γδ T-cells were washed twice with PBS to remove TransAct (a non-polymeric CD3/CD28 activator), then resuspended with neutralized collagen solution to reach a concentration of 5×10^5^ cells/ ml. Crosslinking was done by adding methyl-tetrazine-peg 5-methyl-tetrazine to gels for 20 minutes at room temperature to allow for diffusion, then at 37C for 1 hour to allow the crosslinking reaction to occur. Gels were then washed twice using complete T-cell media. Parallel 2D cultures were maintained in 48 or 12-well MatTek dishes as required.

#### Isolation from culture matrix

To isolate γδ T cells from the culture matrix, Col-nb gels were first transferred into a new sterile plate with fresh complete T-cell media to ensure only encapsulated cells were collected. Gels were then mechanically dissociated, followed by enzymatic digestion, using a collagenase type-I solution.

### Cancer cell culture

Luciferized Human colorectal carcinoma cells (HCT116) received as a gift from the Ingber lab, were cultured with RPMI 1640(Lonza, BE12-702F) with 10% heat inactivated fetal bovine serum (Gibco,10-082-147) and 1% penicillin/ streptomycin. HCT 116 cells used for the intracellular cytokine stimulation assay, were passaged at least 6 times. 4×10^3^ HCT116 cells were seeded in 96-well tissue culture plastic plates overnight to allow for adhesion. γδ T-cells were then co-cultured at a 3:1 ratio for 4 hours.

### Flow cytometry

γδ T-cells were kept at 4°C throughout immunostaining. Cells were stained with near IR live dead cell stain (Fisher Scientific, L10119) according to the manufacturer’s protocol. Cells were then blocked with FcX Fc receptor blocking solution (Biolegend, 422302) for 20 min and stained with surface protein antibodies for 30 min. Brilliant violet staining buffer (BD Biosciences, 566349) and flow cytometry staining buffer (eBioscience, 00-4222-26) were used during staining. For intracellular staining, after stimulation with HCT116 cancer cells for 1hr, Golgi plug (BD Biosciences, BDB555029) was added, and after 4 hours samples were transferred to u-bottom 96-well plates. Fixing and permeabilization were carried out using the CytoFast fix and perm kit (Biolegend, NC1666067). Intracellular staining was performed per manufacturer’s instructions. Flow cytometry acquisition was carried out with the Sony ID7000 spectral cytometer. Gating was performed based on fluorescence-minus-one controls after compensation was adjusted as recommended by Sony (Fig.S4). The complete set of antibodies used for flow cytometry are listed in Supplementary Table (1)

### Proteomics

#### Sample preparation for proteomics

Following retrieval from the collagen matrices, γδ T-cells were washed and stained with Near-IR Live/Dead stain (Fisher Scientific, L10119) and antibodies against CD3, Vδ1, and Vδ2. The stained cells were then sorted by FACS and frozen in PBS prior to proteomic sample preparation.

#### Protein Extraction, Reduction and Alkylation Protocol

The sorted cells were thawed and lysed in 6M Guanidinium Chloride (spec) (1:1 v/v) and vortexed for 2 minutes. 10uL of 10mM TCEP (tris(2-carboxyethyl)phosphine) in 50mM Tetraethylammonium bicarbonate (TEAB) was added to each sample, followed by incubation for 45 minutes at 25°C while shaking at 350rpm using an Eppendorf ThermoMixer C. Following reduction, 10uL of 10mM iodoacetamide in 50mM TEAB was added to the samples for alkylation, with the samples incubated for 15 minutes at 25°C while shaking at 350 rpm.

#### SP3 Magnetic Bead Digestion

Following reduction and alkylation, the extracted protein samples (10 μg per sample) were processed using a modified SP3 (Single-Pot Solid-Phase-enhanced Sample Preparation) protocol to enable efficient protein binding, clean-up, and on-bead digestion.

#### Bead Preparation

Magnetic beads (Promega, cat. number CS3325A04) were vortexed thoroughly to ensure homogeneity, and 30 μL (500 μg) aliquots were transferred for each sample, in accordance with the vendor’s protocol. Beads were washed three times with high-purity water using a magnetic rack to separate beads and supernatant discarded between washes.

#### Protein Binding

Reduced and alkylated protein samples were added to the washed beads. Ethanol was added to achieve an 80% final ethanol concentration. The mixture was vortexed for 30 seconds and incubated with agitation at 1,200 rpm for 20 minutes to promote protein binding to the beads. Beads were then captured on a magnetic rack, supernatants removed, and the bead-protein complexes were washed three times with 1 mL of 80% ethanol.

#### On-Bead Digestion

Proteins bound to beads were resuspended in 50 mM TEAB buffer and Trypsin Platinum protease MS grade (Promega, Ref VA900A) (100mg/mL) was added at a 1:50 enzyme-to-protein ratio to the samples. Digestion proceeded overnight at 40°C with agitation at 1,200 rpm. Post-digestion, beads were separated magnetically and the supernatant collected. Beads were then washed once with 50 mM TEAB for maximum collection of peptides, and the wash combined with the initial supernatant. The final peptide solution was dried and resuspended in 10 μL of Buffer A (0.1% formic acid in ultrapure HPLC grade water) for subsequent LC-MS/MS analysis.

#### Mass spectrometry analysis

Each sample was then submitted for single LC-MS/MS experiment that was performed on a Orbitrap Astral (Thermo Scientific, MA) equipped with NEO nano pump ( Thermo Scientific, MA). Peptides were separated onto a PepSep C18 15cmx75um, 1.9um analytical column (Bruker Daltonics, MA). Separation was achieved through a 24 min active reverse phase gradient with a 30 min total cycle time. Mobile phase A was 0.1% formic acid in water and mobile phase B was 0.1% formic acid in acetonitrile, and the flow rate was 250 nL/min. Electrospray ionization was enabled through applying a voltage of 2 kV using a PepSep electrode junction at the end of the analytical column and sprayed from stainless still PepSep emitter SS 30µm LJ (Bruker, MA). The Astral Orbitrap was acquired in data-independent acquisition mode for the mass spectrometry methods. The mass spectrometry MS1 survey scan was performed across the scan range of 400 –800 m/z at a resolution of 240K, followed by DIA scans with isolation window of 2 Da. The AGC was set to 200%, the maximum ion accumulation time was set to 20 ms, normalized collision energy was set to 25V.

#### Proteome data analysis

The raw data was submitted for analysis by PEAKS 13 (BSI, Toronto, Canada). The assignment of MS/MS spectra was performed by searching the data against a protein sequence database including all entries from the Human Uniprot database (Homo Sapiens Human Uniprot) and other known contaminants such as human keratins and common lab contaminants. Searches were performed using a 10 ppm precursor ion tolerance and requiring each peptide’s N-/C-termini to adhere with Trypsin protease specificity, while allowing up to two missed cleavages. The static modifications and carbamidomethyl on cysteine amino acids (+57.021464 Da) while methionine oxidation (+15.99492 Da) was set as a variable modification. A MS2 spectra assignment false discovery rate (FDR) of 1% on protein level was achieved by applying the target-decoy database search. Filtering was performed using a Percolator (64bit version, reference 1). For quantification all numbers were automatically assigned by PEAKS software of PSM and peptide levels. Al PSM data were used to do quantitative analysis based on in house made WASP software.

#### Database Search and Bioinformatics Differential Expression

Peptide Spectrum Matches (PSMs) were aggregated to the peptide level via PEAKS (version 13). Next, peptide level abundances were normalized using the Trimmed Mean of M values (TMM) method, null values imputed by left-tail sampling, log_2_ transformed, and aggregated to the protein level by the mean of the top 3 most intense peptides.

#### Differential abundance and visualization

Protein abundances were analyzed from an aggregated, imputed protein matrix in log_2_ space. For each contrast (Slow vs Fast, Fast vs 2D), we performed donor-matched differential analysis using paired measurements: for each protein, log_2_ fold-change was computed as the mean within-donor difference between conditions (A − B) across donors with available values, and statistical significance was assessed using a paired two-sided t-test across donors. Multiple-testing correction across proteins within each contrast was performed using the Benjamini–Hochberg false discovery rate (FDR). Volcano plots display log_2_ fold-change versus −log_10_(raw p-value); points exceeding predefined thresholds (|log_2_FC| ≥ 1 and p < 0.05) were classified as significantly increased or decreased. Preselected proteins of interest were annotated on volcano plots

#### Heatmap visualization

For each contrast, a heatmaps were generated to visualize expression patterns of the most informative proteins across paired samples. Proteins were ranked using a composite score defined as (|log_2_FC| x -log_10_(p)), where (log_2_FC) and (p) are from the paired differential analysis. The top 30 proteins were selected. For visualization, only donors present in both conditions were included, and columns were ordered in an interleaved paired structure, to preserve within-donor comparisons. Values were standardized per protein by row-wise z-scoring across the displayed samples (mean-centered and scaled by the row standard deviation; ddof=1) and clipped to the range [−2.5, 2.5].

#### Pathway enrichment

Over-representation analysis (ORA) was carried out separately on significantly up- and down-regulated gene sets (defined by fold-change and p-value thresholds) using Enrichr libraries specified a priori. Enrichment significance was reported using Enrichr-adjusted p-values, and results were visualized as diverging bar plots showing ±−log_10_ (adjusted p-value) for the top enriched terms per direction and library. In addition, preranked gene set enrichment analysis (GSEA) was performed using a signed ranking metric defined as sign(log_2_FC) × −log_10_(p-value) to capture both effect direction and significance; enrichment was evaluated against selected pathway libraries and summarized using standard GSEA outputs.

### Cell motility and migration

#### Confocal Imaging

Confocal microscopy imaging of the Col-nb gels with encapsulated cells were performed on a temperature and CO_2_ controlled microscope (STELLARIS Leica). Images were scanned in two dimensions along the y and x axis at different z levels using a bidirectional laser. Tiles and z levels were selected based on the highest observed cell density. The total number of tiles selected per time point was twenty-four and each tile was imaged through 180 cycles. Cells were imaged for 6 hours and 30 minutes in 2 minutes and 18 second intervals, which was the minimum time needed to complete 24 tiles for each cycle. The Col-nb gels were imaged on Day 1 and Day 5.

#### Image Analysis

The migratory analysis was performed by Trackmate, a detector interface distributed by Fiji^69^. Specifically, this detector applies a LoG (Laplacian of Gaussian) filter to the images in either SIR-DNA or TRANS-TMPT channel. The calculations were made in the Fourier space. The local maximums in the filtered image are searched for, and maximums that are too close are suppressed, creating an almost binary image that allows for subpixel localization. We selected for cells roughly 10 microns in diameter. We selected the linking maximum distance and a gap-closing max frame gap to be 15-30 microns. Trackmate helped calculate the distance traveled between each frame for each cell, which is marked as a “spot”.

### Xenograft HCT116 model

#### Tumor inoculation and γδ T-cell treatment

Animal studies were approved by the National Institutes of Health and the Harvard University Faculty of Arts and Sciences Institutional Animal Care and Use Committee (IACUC).

6-week-old Female NSG human IL-15 producing mice obtained from the Jackson laboratory (030890) were inoculated with 2.5×10^5^ HCT116 luciferized cells subcutaneously. After 7 days, tumor bearing mice were injected subcutaneously with 5×10^4^ γδ T-cells from either slow-relaxing, fast-relaxing, or 2D conditions. Saline only (non-treated mice) were used as the negative control group.

#### Tumor tracking

Tumor progression was monitored via luminescence imaging and caliper measurements. Animals were anesthesized and peritumorally injected with D-luciferin (ThermoFisher L2916), at 150mg kg^-^.Luminescence was measured over 30 minutes with 5 minute intervals, using IVIS. Mice tumor progression measurements were carried out every two days, with euthanasia performed after tumors reached a size of 2,000mm^3^ or became ulcerated.

### Statistical analyses

Unless otherwise specified in figure legends, statistical analyses were performed on Prism GraphPad. Comparisons were performed using two-tailed one-way paired ANOVA, and post-hoc tests for multiple comparisons, or Student’s t-test or Wilcoxon test for comparison between two groups. Prism mixed-effects model analysis was used for datasets with missing values (where ordinary two-way ANOVA could not be performed). P < 0.05 was considered significant unless otherwise noted. Error bars represent the standard deviation of the mean, unless otherwise noted.

## Data Availability

The authors declare that all data supporting the findings of this study are available within the paper and its supplementary information files. The RNA-sequencing datasets analyzed in this study, were assessed from publicly available data with accession codes GSE263248, and GSE164378.

## Acknowledgements

This work was supported by the National Institute of Health/ National Cancer Institute grant no. 5R01CA276459. F.O acknowledges funding from the National Science Foundation Graduate Research Fellowship, the Ford Foundation Fellowship and the MIT University Center for Exemplary Mentoring Fellowship. We thank Drs. Kwasi Adu Berchie, Yoav Binenbaum, Wei-Hung Jung, Nuria Lafuente Gomez and Shawn Kang for their scientific inputs.

## Author contributions

Conceptualization: F.O, D.J.M, Methodology: F.O Investigation: F.O, M.C, J.P, T.T, L.L, Proteomics: F.O, B.B, Visualization: F.O, M.C, J.P, Funding acquisition: F.O, D.J.M, Writing – original draft: F.O, Writing – review & editing: F.O, M.C, J.P, D.J.M

