## Supplementary figure (S) for "Viscoelastic Niches Shape γδ T-Cell Phenotype and Effector Function"

### Supplementary Figures and Tables

Favour Omafuvwe Obuseh [1,3], Michelle Chang [2], Joshua Price [2,3], Tania To [2,3], Kyle Ruark[2,3],Leah Lourenco[2], Bogdan Budnik[3], David Mooney[2,3]

**Affiliations:**

1. Harvard-MIT Program in Health Sciences and Technology (HST), Cambridge, MA, USA
2. School of Engineering and Applied Sciences (SEAS), Harvard University, Cambridge, MA, USA
3. Wyss Institute for Biologically Inspired Engineering, Harvard University, Boston, MA, USA

### Supplementary Figures

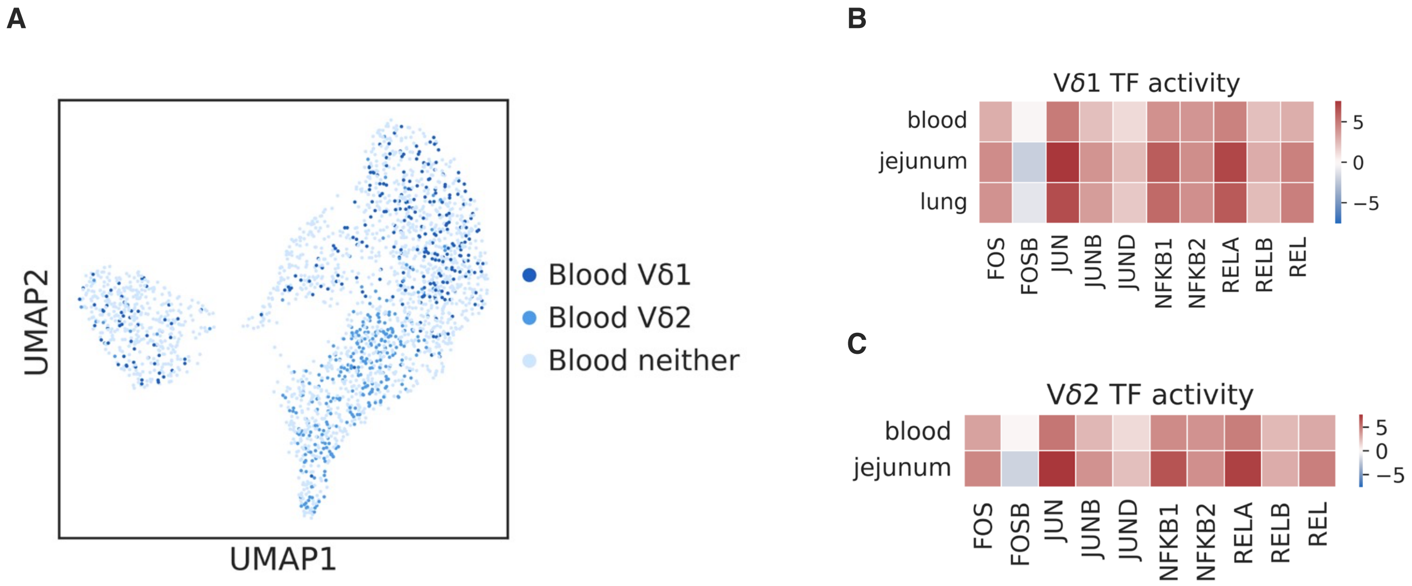


**Figure S1**. **Single cell RNA-sequencing reveals tissue-dependent transcriptional states in primary human γδ T-cells**
(A) UMAP of γδ T-cells from blood, indicating Vδ1 and Vδ2 identities (sequencing data from Hao et al., 2021). (B) CollecTRI ULM t-values for FOS, JUN, and NFKB family transcription factors in blood, jejunum, and lung Vδ1 cells; mean across donors. (C) CollecTRI ULM t-values for FOS, JUN, and NFKB family transcription factors in blood and jejunum Vδ2 cells; mean across donors. B represent 4 jejunum, 2 lung, and 6 blood donors, C represent 4 jejunum, and 6 blood donors

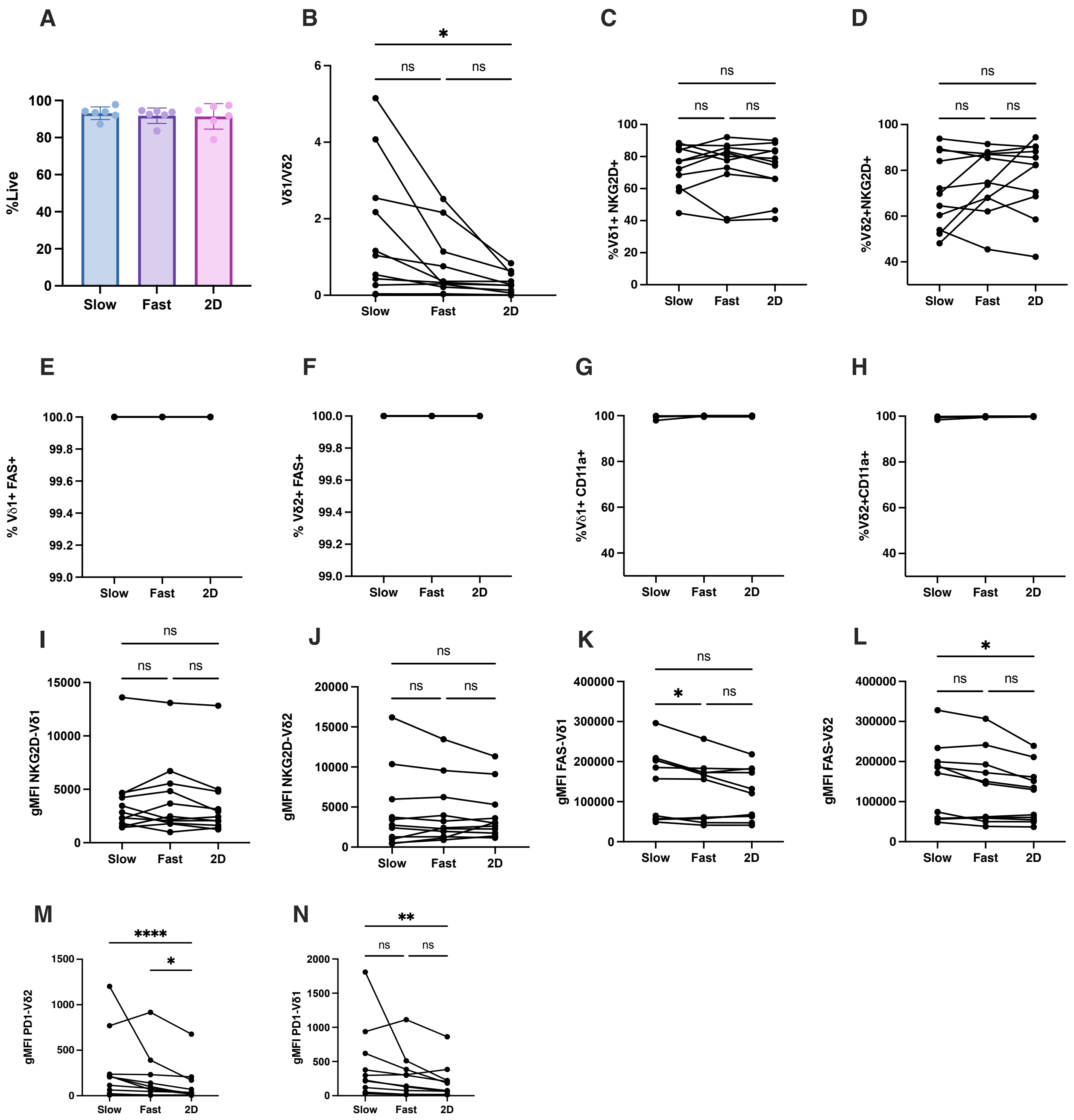


**Figure S2. Viscoelasticity Effect on Activation**

(A) Percentage of viable γδ T-cells after encapsulation in slow-relaxing, fast-relaxing, or unencapsulated (2D) conditions. (B) Vδ1/Vδ2 ratio after encapsulation in slow-relaxing, fast-relaxing, or unencapsulated (2D) conditions.(C) Percentage of $V\delta1^+$ $NKG2D^+$ cells after encapsulation in slow-relaxing, fast-relaxing, or unencapsulated (2D) conditions.(D) Percentage of $V\delta2^+$ $NKG2D^+$ cells after encapsulation in slow-relaxing, fast-relaxing, or unencapsulated (2D) conditions.(E) Percentage of $V\delta1^+$ $FAS^+$ cells after encapsulation in slow-relaxing, fast-relaxing, or unencapsulated (2D) conditions.(F) Percentage of $V\delta2^+$ $FAS^+$ cells after encapsulation in slow-relaxing, fast-relaxing, or unencapsulated (2D) conditions.(G) Percentage of $CD11a^+$ Vδ1 cells after encapsulation in slow-relaxing, fast-relaxing, or unencapsulated (2D) conditions.(H) Percentage of $CD11a^+$ Vδ2 cells after encapsulation in slow-relaxing, fast-relaxing, or unencapsulated (2D) conditions.(I) Geometric mean fluorescence intensity (gMFI) of NKG2D in $V\delta1^+$ cells after encapsulation in slow-relaxing, fast-relaxing, or unencapsulated (2D) conditions.(J) Geometric mean fluorescence intensity (gMFI) of NKG2D in $V\delta2^+$ cells after encapsulation in slow-relaxing, fast-relaxing, or unencapsulated (2D) conditions.(K) Geometric mean fluorescence intensity (gMFI) of FAS in $V\delta1^+$ cells after encapsulation in slow-relaxing, fast-relaxing, or unencapsulated (2D) conditions.(L) Geometric mean fluorescence intensity (gMFI) of FAS in $V\delta2^+$ cells after encapsulation in slow-relaxing, fast-relaxing, or unencapsulated (2D) conditions.(M) Geometric mean fluorescence intensity (gMFI) of PD-1 in $V\delta1^+$ cells after encapsulation in slow-relaxing, fast-relaxing, or unencapsulated (2D) conditions.(N) Geometric mean fluorescence intensity (gMFI) of PD-1 in $V\delta2^+$ cells after encapsulation in slow-relaxing, fast-relaxing, or unencapsulated (2D) conditions. For panel A, n = 6 donors. For panel B, n = 11 donors. For panels C–N, n = 11 donors. Bar graphs show mean ± s.d. Statistical significance was determined using one-way ANOVA with Tukey's multiple comparisons test. *$P < 0.05$, **$P < 0.01$, ***$P < 0.001$, ****$P < 0.0001$; ns, not significant.

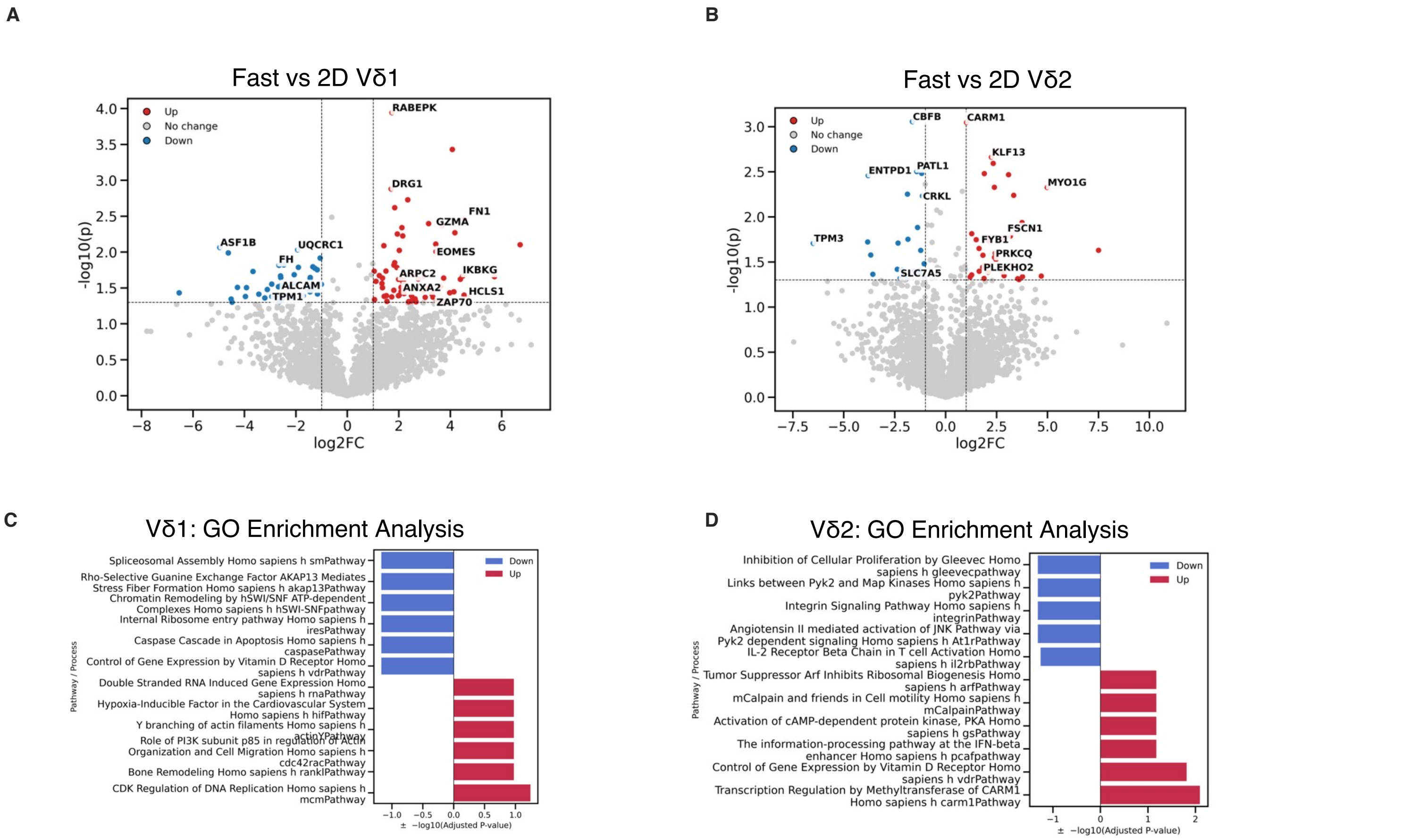


**Figure S3. Encapsulation Drives Proteomic Remodeling in γδ T-cells**

(A) Volcano plot comparing FACS-sorted Vδ1 cells cultured in fast-relaxing matrix conditions versus non-encapsulated. (B) Volcano plot comparing FACS-sorted Vδ2 cells cultured in fast-relaxing matrix conditions versus non-encapsulated conditions. (C) Enrichr over-representation analysis (ORA) of Gene Ontology (GO) biological processes (BioCarta 2016) for Vδ1 cells cultured in fast-relaxing matrix conditions versus non-encapsulated conditions. (D) Enrichr over-representation analysis (ORA) of Gene Ontology (GO) biological processes (BioCarta 2016) for Vδ2 cells cultured in fast-relaxing matrix conditions versus non-encapsulated conditions. For panels A,C, n = 4 donors; for panels B,D, n = 3 donors. Statistical significance in A,B, was determined using a paired t-test $^*P < 0.05$, $^{**}P < 0.01$, $^{***}P < 0.001$; ns, not significant.

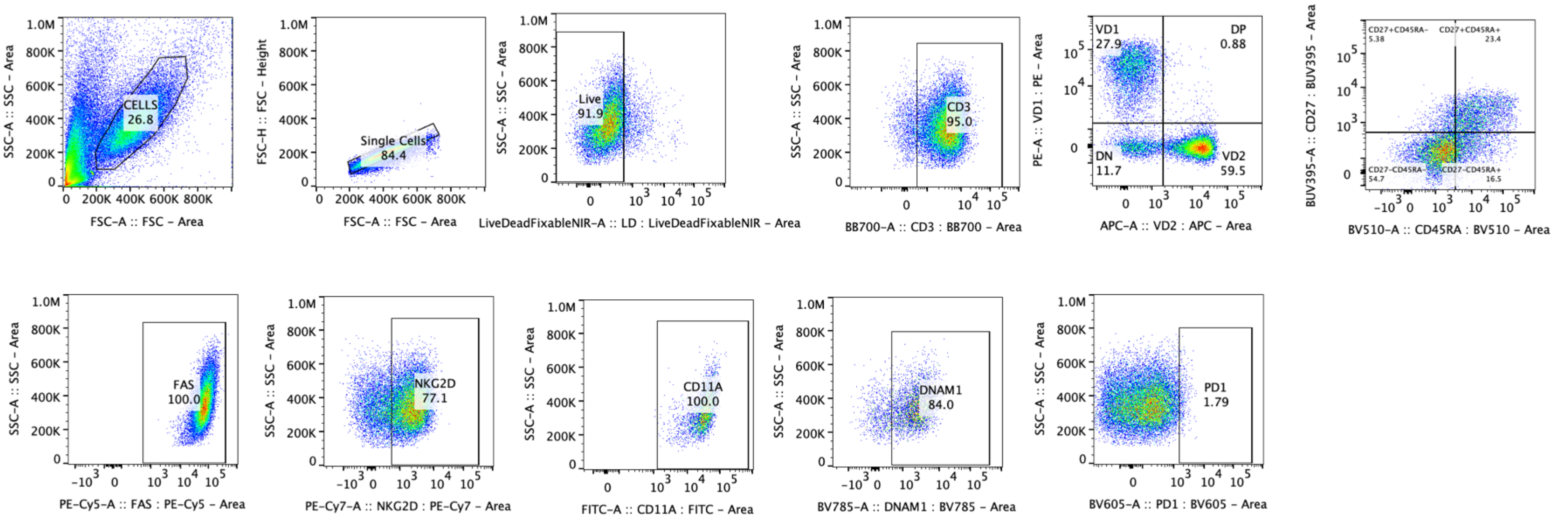


**Figure S4. Gating Strategy**

Gating performed using fluorescence minus one (FMO) control.

| Staining Antibody | Manufacturer | Cat# |
| --- | --- | --- |
| CD3 (RB705) | BD Biosciences | 570237 |
| TCRγδ (PE/Dazzle) | BioLegend | 331226 |
| TCR Vδ1 (PE) | Thermo Fisher | 12-5679-42 |
| TCR Vδ2 (APC) | BioLegend | 331418 |
| CD27 (BUV395) | Thermo Fisher | 363027942 |
| CD45RA (Brilliant Violet 510) | BioLegend | 304142 |
| PD-1 (BV605) | BioLegend | 367426 |
| NKG2D (PE-Cy7) | Thermo Fisher | 25-5878-42 |
| CD11a (FITC) | Thermo Fisher | 11-0119-42 |
| DNAM-1 (BV785) | BioLegend | 338322 |
| NKp30 (BV711) | BioLegend | 325218 |
| CD95/FAS (PE-Cy5) | Thermo Fisher | 15095942 |
| Live/Dead NIR viability dye | Thermo Fisher | L10119 |
| TIGIT (BV421) | BioLegend | 372710 |
| IFNγ (FITC) | BD Biosciences | 554551 |
| Granzyme B (PE-Texas Red) | Invitrogen | GRB17 |
| TNFα (BV421) | Thermo Fisher | 404-7349-42 |

**Table S1. Antibodies used for experiments**

List of antibodies used for flow cytometry, with manufacture's catalog number provided. The antibodies were used at 1:60 dilution, except for Live Dead NIR which was 1:1000 dilution.
